# JNK signaling-mediated exocytosis coordinates epithelial cell-turnover in *Drosophila* ribosomal protein mutants

**DOI:** 10.1101/2023.12.27.573431

**Authors:** Nanami Akai, Tatsushi Igaki, Shizue Ohsawa

## Abstract

Robust tissue growth is orchestrated by the precise coordination of cell death and cell proliferation. Our previous study found that in the developing wing pouches of *Drosophila Minute*/+ animals, both cell death and compensatory cell proliferation are dramatically increased, which contributes to robust growth of mutant tissue. The induction of this cell-turnover depends on activation of JNK signaling, although the mechanism by which JNK activation induces cell-turnover remained unclear. Here, we show that JNK-mediated elevation of exocytosis in dying cells is crucial for triggering cell-turnover in *M/+* wing morphogenesis. Mechanistically, elevated JNK signaling in dying cells upregulates exocytosis-related genes and Wingless (Wg), leading to enhanced Wg secretion. Furthermore, this exocytosis-mediated Wg secretion generally occurs downstream of JNK signaling, regardless of the genetic background. Overall, our findings provide mechanistic insights into robust tissue growth through the orchestration of cell-turnover, which is primarily governed by JNK-mediated exocytosis during *Drosophila Minute/+* wing morphogenesis.

## Introduction

Ribosomes are essential molecular machines responsible for protein synthesis in living cells, and thus fundamental to life. In humans, heterozygosity of genes related to ribosomal proteins or ribosomal biogenesis factors leads to a group of genetic disorders collectively termed ribosomopathies. For instance, heterozygous mutations in various ribosomal protein genes, including *RPS17* ^1^*, RPS26* ^2^, *RPS19* ^3^, *RPS24* ^4^, and *RPL27* ^5^, are associated with Diamond-Blackfan anaemia (DBA), with some patients exhibiting tissue-specific developmental anomalies such as limb defects, cleft palate, and abnormal heart development ^6,7^. In addition, heterozygous mutations in the *RpSA* gene are linked to isolated congenital asplenia (ICA), a disorder characterized by the absence of a spleen at birth ^8^. However, the precise mechanisms by which a reduction in ribosomal protein gene dosage leads to these genetic disorders remain elusive.

In *Drosophila*, a series of heterozygous mutants for ribosomal protein genes, called *Minute/+* (*M/+*), exhibit a pronounced developmental delay during larval stages ^9^. Despite this significant delay, *M*/+ animals are essentially normal flies without noticeable morphological defects, except for the thinner bristles, suggesting that *M/+* animals exert certain mechanisms to overcome developmental perturbations caused by reduced ribosomal protein levels. It has been reported that extensive cell death ^10^, as well as cellular stress ^10–13^, is a distinctive characteristic of the *M*/+ wing pouch. Our previous study ^14^ has shown that apoptotic cell death and the subsequent cell proliferation are dramatically increased in the *M*/+ wing pouch. Blocking this cell-turnover by inhibiting cell death resulted in morphological abnormalities, indicating the essential role of cell-turnover in *M/+* wing morphogenesis. Genetic analyses have revealed that the induction of this cell-turnover depends on activation of JNK signaling. However, downstream events of JNK activation remain to be elucidated.

The coordination of cell death and proliferation through cell-cell communications is crucial for proper development and homeostasis of multicellular organisms. Apoptotic cells, for example, can secrete mitogens such as Wingless (Wg; a Wnt homolog), dpp (a BMP homolog), and Hh, which could promote the proliferation of nearby cells in the *Drosophila* epithelium ^15–19^. Cell competition, a phenomenon in which cells with higher fitness (“winners”) eliminate neighboring less fit cells (“losers”) by inducing cell death ^20^, is another aspect of cell-turnover. It has been reported that secretory factors from prospective loser cells contribute to cell competition. For instance, the cytokine-like ligand unpaired 3 (upd3) produced by losers promotes cell competition between wild-type (winners) and *M/+* cells (losers) in the *Drosophila* epithelium ^12^. Furthermore, Madin-Darby canine kidney cells depleted of the polarity regulator gene *scribble* (*scrib*^KD^ MDCK cells) secrete Fibroblast Growth Factor 21 (FGF21), which in turn promotes cell competition with neighboring wild-type MDCK cells ^21^. However, the mechanism by which these secreted factors are released from dying cells is still not fully understood.

Here, we found that exocytosis, acting as a downstream event of JNK signaling in dying cells, contributes to cell-turnover in the *M/+* wing pouch, which is essential for robust wing development in *M*/+ animals. Our genetic analyses suggest that elevated exocytosis likely facilitates the release of Wg from dying cells, which in turn stimulates their own cell death and the proliferation of neighboring living cells. Furthermore, we found that this exocytosis-mediated Wg release is not limited to *M*/+ context but represents a general event downstream of JNK signaling. Our observations could provide new mechanistic insights into how dying cells coordinate robust development through cell-cell interactions.

## Results

### *M/+* wing pouch elevates exocytosis downstream of JNK signaling

To elucidate the downstream event of JNK signaling in the *M/+* wing pouch, we conducted RNA sequence (RNA-seq) analyses on GFP-labeled fluorescence-activated cell sorting (FACS)-sorted wing pouch cells from one of the *M/+* mutants *RpS3/+*, compared to GFP-labeled wild-type control or *RpS3/+* wing pouch cells overexpressing JNK inhibitor Puckered (Puc) (Figure S1A; the wing pouch is green-marked oval domain that becomes the adult wing blade). Among the 1,097 genes that were upregulated or downregulated in the *RpS3/+* wing pouch relative to the wild-type control, we identified positive regulators of JNK signaling such as *reaper* and *pvf1*, JAK/STAT targets including *Socs36E* and c*hinmo*, Gr64 gustatory receptors and numerous genes associated with oxidative stress and DNA repair (Figure S1B, Tables S1 and S2), which aligns with previous reports on genes upregulated in the *RpS3/+* or *RpS17/+* wing disc ^12,13^. These data validate our experimental conditions for identifying genes required for JNK-mediated events in the *M/+* wing pouch.

Mining the list of genes differentially expressed in the *RpS3/+* wing pouch cells dependent on JNK signaling (Figure S1C and Table S3), we noticed that among the genes associated with the “secretion by cell” GO term (Figure S1B), the evolutionarily conserved exocytosis-related genes *unc-13*, *SNAP25,* and *cadps* (*Calcium-dependent secretion activator*) were upregulated in a JNK-dependent manner (2.29-fold increase, *RpS3*/+ compared to wild-type; 3.44-fold increase, *RpS3*/+ compared to *RpS3*/+ + Puc) (Figure S1D). Among these exocytosis-related genes, the upregulation of *SNAP25* and *unc-13* was consistent with the previous reports concerning genes upregulated in *RpS3/+* or *RpS17/+* cells ^12,13^ .

Calcium-dependent exocytosis is the process whereby cells release intracellular contents into the extracellular space. It has been shown that *unc-13, SNAP25*, and *cadps* collectively regulate the docking process of secretory vesicles to the plasma membrane during Ca^2+^-mediated exocytosis ^22^. Specifically, UNC-13, a conserved presynaptic protein with calcium-binding domains, interacts with syntaxin to prime vesicles for fusion, crucial for calcium-regulated exocytosis ^23^. SNAP-25, in conjunction with syntaxin-1 and synaptobrevin, forms the pivotal SNARE complex for neuronal exocytosis, assembling into a four-helix bundle that is essential for drawing vesicle and plasma membranes close together to enable membrane fusion ^24,25^. Like UNC-13, CAPS/Cadps possesses conserved C-terminal domains that are instrumental in the assembly of SNARE complexes, thus priming vesicles for Ca^2+^-induced exocytosis ^26^. First, to investigate whether exocytosis is upregulated in the *M/+* wing pouch, we utilized the exosome marker CD63 tagged with EGFP (EGFP-CD63) ^27–30^. Additionally, we utilized Syt1 tagged with EGFP (Syt1-EGFP), a widely used vesicle marker prevalent in both neuronal and non-neuronal cells ^31–33^. We observed a significant increase in EGFP-CD63-positive and Syt1-EGFP-positive vesicles in the *RpS3/+* wing pouch compared to the wild-type control (Figures 1A, 1B, 1G and 1H, corresponding to the boxed region in Figure 1E, quantified in Figures 1F and 1K). EGFP-CD63-positive vesicles were also found to be increased in the *RpL19/+* wing pouch (Figure S1E, quantified in Figure S1F). Notably, these vesicles were predominantly observed in regions of the *RpS3/+* wing pouch undergoing massive cell death (as indicated by arrowheads in Figures 1B, 1H and S1E), a phenomenon we previously reported ^14^ and also confirmed in this study (shown in Figure 2H, compared to Figure 2G, quantified in Figure 2T). Furthermore, we noted an increase in vesicles positive for the ESCRT protein Hrs, an additional exosome marker crucial for exosome secretion ^34,35^, in the *RpS3/+* wing pouch relative to the wild-type control (Figures S1G and S1H, quantified in Figure S1I). To further investigate the enhanced exocytosis in the *RpS3/+* wing pouch, we utilized the GCaMP6m fluorescent calcium reporter ^36^ to monitor Ca^2+^ activity, a known inducer of exocytosis. We observed persistent Ca^2+^ activity in morphologically dying cells, along with spontaneous Ca^2+^ flashes similar to those in the wild-type control (Figures 1L and 1M, quantified in Figure 1P). This persistent activity suggested an enhanced exocytic process in dying cells in the *M/+* wing pouch, potentially driven by elevated calcium levels.

**Figure 1.**
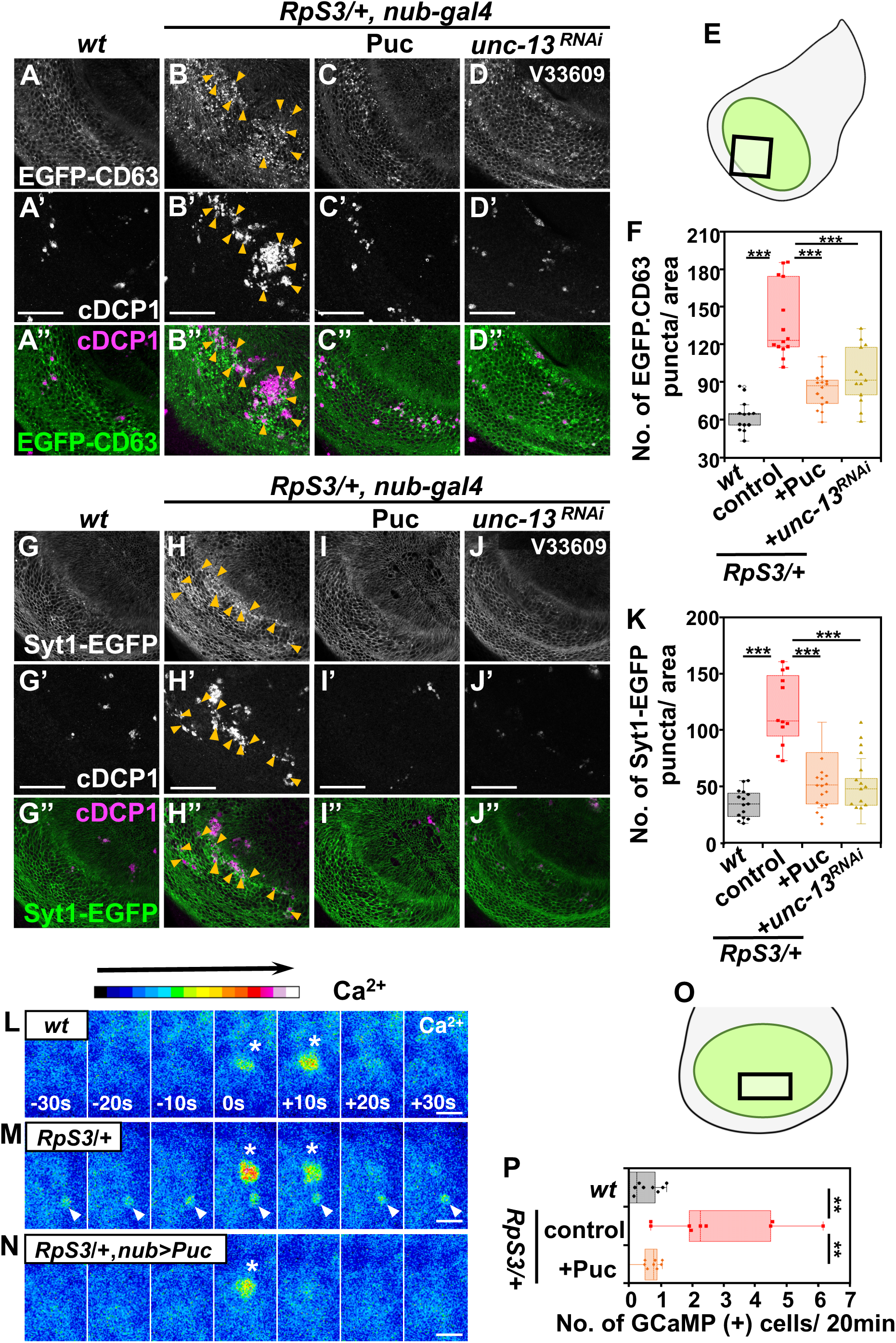
*M/+* wing pouch elevates exocytosis downstream of JNK signaling. **(A-D’’)** The exosome marker EGFP-CD63 was expressed in the wing pouch of wild-type (A), *RpS3/+* (B), *RpS3/+, nub-Gal4, UAS-Puc* (C), or *RpS3/+, nub-Gal4, UAS-unc-13-RNAi* (D) flies (white). The images correspond to the boxed area in the schematic diagram of the wing disc (E). Dying cells in the wing disc were visualized by anti-cleaved Dcp-1 staining (white). EGFP-CD63-positive vesicles were frequently observed in the area of morphological dying cells in the *RpS3/+* wing pouch (Indicated by Orange arrowheads). Scale bar, 50 μm. **(F)** Boxplot with dots representing the number of EGFP-CD63-positive puncta in the pouch of genotypes shown in (A) (n=13, number of wing pouches), (B) (n=14), (C) (n=16), and (D) (n=13). Error bars, SEM; ***, p<0.001; Wilcoxon rank-sum test. **(G-J’’)** The vesicle marker Syt1-EGFP was expressed in the wing pouch of wild-type (G), *RpS3/+* (H), *RpS3/+, nub-Gal4, UAS-Puc* (I), or *RpS3/+, nub-Gal4, UAS-unc-13-RNAi* (J) flies (white). The images correspond to the boxed area in the schematic diagram of the wing disc (E). Dying cells in the wing disc were visualized by anti-cleaved Dcp-1 staining (white). Syt1-EGFP-positive vesicles were frequently observed in the area of morphological dying cells in the *RpS3/+* wing pouch (Indicated by Orange arrowheads). Scale bar, 50 μm. **(K)** Boxplot with dots representing the number of Syt1-EGFP-positive puncta in the wing pouch of genotypes shown in (G) (n=15), (H) (n=12), (I) (n=14), and (J) (n=16). Error bars, SEM; ***, p<0.001; Wilcoxon rank-sum test. **(L-N)** Time-lapse imaging of Ca^2+^ signaling in cultured wing discs. The calcium reporter GCaMP6m (shown in pseudo color) was expressed in the wing pouch of wild-type (L), *RpS3/+* (M), or *RpS3/+, nub-Gal4, UAS-Puc* (N) flies. Images were acquired at 10-second intervals. White arrowheads indicate the persistent activation of Ca^2+^ signaling. White asterisks indicate spontaneous Ca^2+^ flashes. The images correspond to the boxed area in the schematic diagram of the wing disc (O). Scale bar, 5 μm. **(P)** Quantification of cells exhibiting continuous Ca^2+^ activity for 20 min in the wing pouch. The respective genotypes are shown in (L) (n=12), (M) (n=10), and (N) (n=9). These images are Z-stacked images from 3 confocal images (4μm). Error bars, SEM; **, p<0.01; Wilcoxon rank-sum test.

**Figure 2.**
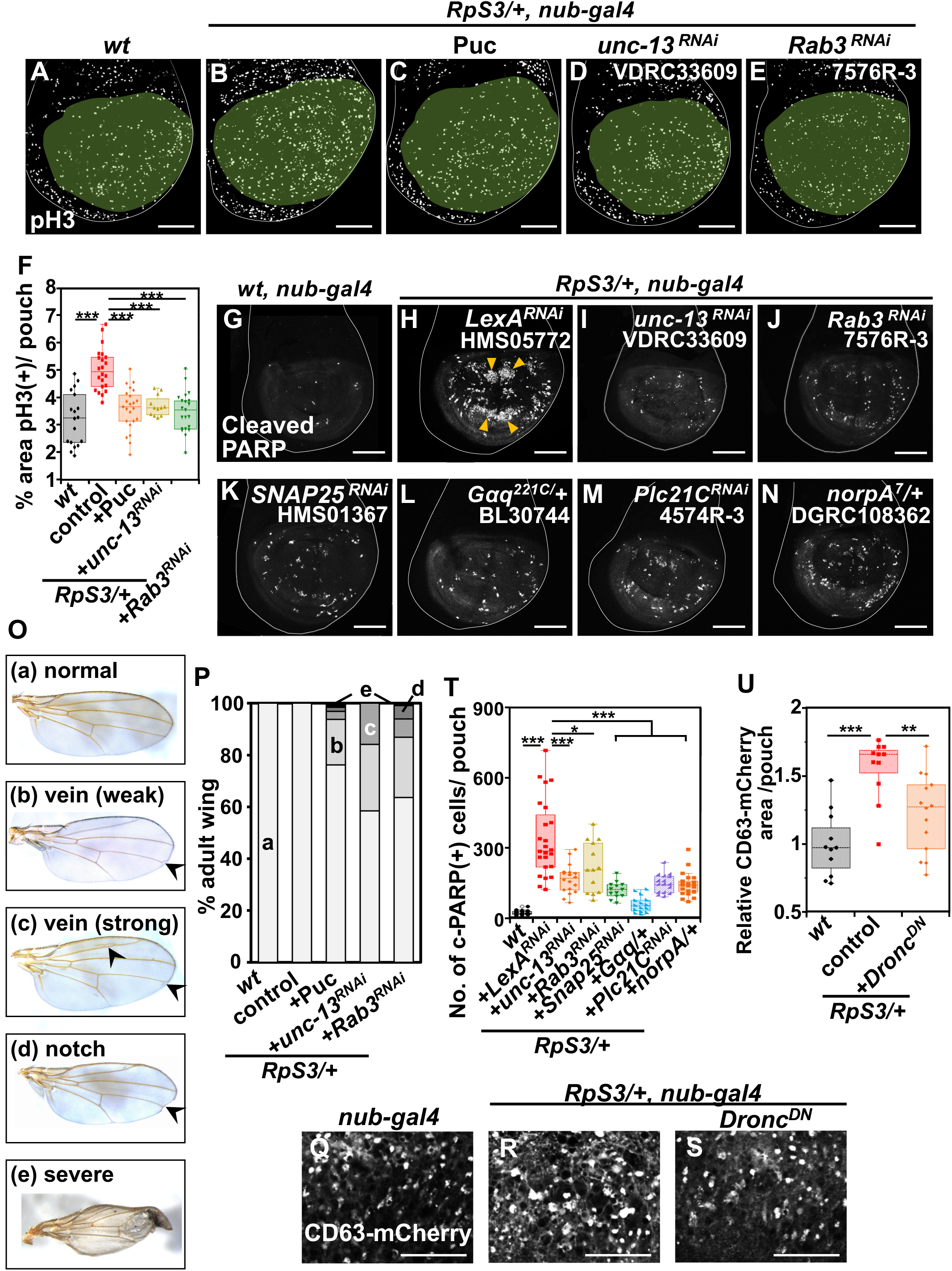
Exocytosis is required for massive cell-turnover in the *M/+* wing pouch. **(A-E)** Wing discs of wild-type (A), *RpS3/+* (B), *RpS3/+,*), *RpS3/+, nub-Gal4, UAS-unc-13-RNAi* (D), or *RpS3/+, nub-Gal4, UAS-Rab3-RNAi* (E) flies were stained with anti-phospho-histone H3 (pH3) (Ser10) antibody (white). Wing pouches were marked in pale green. Scale bar, 100 μm. **(F)** Boxplot with individual dots representing pH3-positive areas in the pouch of genotypes shown in (A) (n=18), (B) (n=22), (C) (n=23), (D) (n=12), and (E) (n=21). Error bars, SEM; ***, p<0.001; Wilcoxon rank-sum test. **(G-N)** The activated-caspase-3 indicator CD8-PARP-Venus was expressed in the wing pouch of wild-type (G), *RpS3/+, nub-Gal4, UAS-lexA-RNAi* (H), *RpS3/+, nub-Gal4, UAS-unc-13-RNAi* (I), *RpS3/+, nub-Gal4, UAS-Rab3-RNAi* (J), *RpS3/+, nub-Gal4, UAS-SNAP25-RNAi* (K), *RpS3/+, Gαq/+* (L), *RpS3/+, nub-Gal4, UAS-Plc21C-RNAi* (M), or *RpS3/+, norpA/+* (N) flies, and dying cells in the wing pouch were visualized by anti-cleaved PARP staining (white). Orange arrowheads indicate massive cell death in the *RpS3/+* wing pouch. Scale bar, 100μm. **(O)** Adult wing phenotypes were classified as following five types: (a) normal, (b) vein (weak) (weakly bearing a brunched vein in the point of “landmark 14” (arrow) ^68^ (c) vein (strong) (bearing other additional vein phenotypes (arrowheads) in addition to “landmark 14”), (d) notch, and (e) severe. **(P)** The rate of defective wings in the genotypes of wild-type (n=55), *RpS3/*+ (n=44), *RpS3/+, nub-Gal4, UAS-Puc* (n=63), *RpS3/+, nub-Gal4, UAS-unc-13-RNAi* (n=101), and *RpS3/+, nub-Gal4, UAS-Rab3-RNAi* (n=99). **(Q-S)** The exosome marker CD63-mCherry was expressed in the wing pouch of wild-type (Q), *RpS3/+* (R), or *RpS3/+, nub-Gal4, UAS-Dronc^DN^* (S) flies (white). The images correspond to the area enclosed by rectangle 1 in the schematic diagram of the wing disc (Figure 3C). Scale bar, 20 μm. **(T)** Boxplot with individual dots representing the number of cleaved-PARP-positive dying cells per pouch in genotypes shown in (G) (n=18), (H) (n=24), (I) (n=18), (J) (n=13), (K) (n=13), (L) (n=15), (M) (n=16), and (N) (n=22). Error bars, SEM; ***, p<0.001 , **, p<0.01, *, p<0.05; Wilcoxon rank-sum test. **(U)** Boxplot with dots representing the number of CD63-mCherry-positive puncta in the pouch of genotypes shown in (Q) (n=11, number of wing pouches), (R) (n=11), and (S) (n=14). Error bars, SEM; ***, p<0.001, **, p<0.01; Wilcoxon rank-sum test.

We then examined whether exocytosis is the downstream event of JNK activation in the *M/+* wing pouch. We found that blocking JNK signaling by overexpressing Puc significantly suppressed the increased number of EGFP-CD63/Syt1-EGFP-positive vesicles, as well as the emergence of cells exhibiting persistent Ca^2+^ signaling (Figure 1C, quantified in Figure 1F; Figure 1I, quantified in Figure 1K; Figure 1N, quantified in Figure 1P). Together, these observations suggest that exocytosis is elevated in dying cells within the *M/+* wing pouch as a downstream event of JNK signaling.

### Exocytosis is required for massive cell-turnover in the *M/+* wing pouch

We have previously demonstrated that JNK signaling is required for cell-turnover in the *M/+* wing pouch ^14^. Blocking JNK signaling by overexpressing Puc significantly reduced cell death and mitoses in the *RpS3*/+ wing pouch, as assayed by the CD8-PARP-Venus probe for caspase activity in the imaginal disc ^37–44^, and the M phase marker phospho-Histone H3 (Figures 2A-2C, quantified in Figure 2F; Figures 2G, 2H and S2A, quantified in Figure S2N). In addition, we intriguingly found that downregulating *unc-13*, a docking factor involved in exocytosis, similarly diminished cell death and mitoses (Figures 2I and S2B, quantified in Figures 2T and S2N, respectively; Figure 2D, quantified in Figure 2F), and also reduced EGFP-CD63-positive and Syt1-EGFP-positive vesicles in the *RpS3*/+ wing pouch (Figures 1D and 1J, quantified in Figures 1F and 1K, respectively). Knocking down *rab3*, a “Secretory GTPase Rabs” required for vesicle exocytosis, also significantly reduced cell death and mitoses in the *RpS3*/+ wing pouch (Figures 2J and S2C, quantified in Figures 2T and S2N, respectively; Figure 2E, quantified in Figure 2F). Furthermore, overexpressing Puc or downregulating either *unc-13* or *rab3* led to morphological defects in the *RpS3/+* adult wing, while their individual overexpression or downregulation did not affect wing development (Figures 2O, 2P and S2O). These results suggest that exocytosis, as a downstream component of JNK signaling, is pivotal for cell turnover and subsequent normal morphogenesis in the *M/+* wing pouch. Indeed, downregulating other docking factors such as *SNAP25* or *Syt1* ^45^, another Secretory GTPase *Rab27*, or *ALIX*, an ESCRT (endosomal sorting complexes required for transport)-related protein involved in the biogenesis of extracellular vesicles ^46,47^, also significantly suppressed cell death in the *RpS3*/+ wing pouch (Figure 2K, quantified in Figure 2T; Figures S2D-S2J, quantified in Figure S2N). Additionally, downregulating factors such as *Gαq*, *Plc21C*, and *norpA,* which are required for intracellular Ca^2+^ release that triggers exocytosis process ^32^, also significantly suppressed cell death in the *RpS3*/+ wing pouch (Figures 2L-2N, quantified in Figure 2T; Figures S2K-S2M, quantified in Figure S2N). Together, these data reinforce the vital role of exocytosis in cell turnover within the M/+ wing pouch, functioning as a downstream process of JNK signaling. Intriguingly, blocking Dronc activity by overexpressing Dronc^DN^ suppressed the elevation of exocytosis in the *RpS3*/+ wing pouch (Figure 2S, compared to Figure 2R, quantified in Figure 2U). This suggests the presence of a signal amplification loop between exocytosis and Dronc within the JNK signaling pathway, consistent with previous studies showing Dronc-JNK amplification loop that induce cell death ^48,49^.

### Dying cells secrete Wg via exocytosis in the *M/+* wing pouch

We next sought to identify the protein secreted through exocytosis from dying cells in the *M/+* wing pouch. Our previous genetic analyses ^14^ suggested that cell-turnover in the *M/+* wing pouch is initiated by a mechanism akin to Wg-dependent cell competition, a phenomenon by which cells with higher Wg signaling activity eliminate neighboring cells with lower Wg signaling activity in the wing disc ^50^. Indeed, as demonstrated through previous genetic manipulations ^14^, and confirmed here, genetically reducing the Wg signaling gradient in the entire *M/+* pouch significantly suppressed cell death (Figure S3A, quantified in Figure S3B). Additionally, it has been shown that apoptotic cells produce Wg through JNK signaling ^18,51,52^. These observations led us to hypothesize that dying cells in the *M/+* wing pouch might release Wg via JNK-mediated exocytosis, potentially conferring a survival advantage to their neighboring living cells. Supporting this hypothesis, it has been reported that Wnt/Wg can be released through extracellular vesicles including exosome ^28,53–56^. We found that GFP-Wg-positive puncta, derived from a knock-in allele ^34^, were more abundant in the *RpS3/+* wing pouch compared to the wild-type control (Figures 3A and 3B). This increase in GFP-Wg-positive puncta, particularly in the area with massive cell death within the *RpS3*/+ wing pouch (Figures S3D-S3E’’’, compared to Figure S3C), was more evident when using a membrane-tethered anti-GFP nanobody (Vhh4-CD8), which immobilizes GFP-Wg on the cell surface ^34^ (Figures 3F and 3G, quantified in Figure 3J). Intriguingly, these GFP-Wg-positive puncta, observed both without and with the Vhh4-CD8 morphotrap, sometimes colocalized with CD63-mCherry-positive vesicles (Figures 3E and S3G, indicated by arrowheads), suggesting that GFP-Wg might be secreted via extracellular vesicles through JNK-mediated exocytosis. Supporting this possibility, we found that blocking JNK signaling by overexpressing Puc or knocking down *unc-13* significantly reduced the number of GFP-Wg-positive puncta in the *RpS3/+* wing pouch (Figures 3H and 3I, quantified in Figure 3J). In addition, we observed a significant increase in extracellular GFP-positive puncta and extracellular CD63-mCherry-positive vesicles in the *RpS3/+* wing pouch when GFP-Wg was captured on the cell surface using the Vhh4-CD8 morphotrap, compared to the wild-type control (Figures S3H and S3I, quantified in Figures S3J and S3K). Furthermore, these extracellular GFP-Wg-positive puncta occasionally colocalized with extracellular CD63-mCherry-positive vesicles (6.88% in the wild-type, and 16.45% in the *RpS3/+* wing pouch region, characterized by massive cell death, as indicated by the arrowheads in Figures S3H and S3I). Together, these data suggest that dying cells secrete Wg via exocytosis in the *M/+* wing pouch.

**Figure 3.**
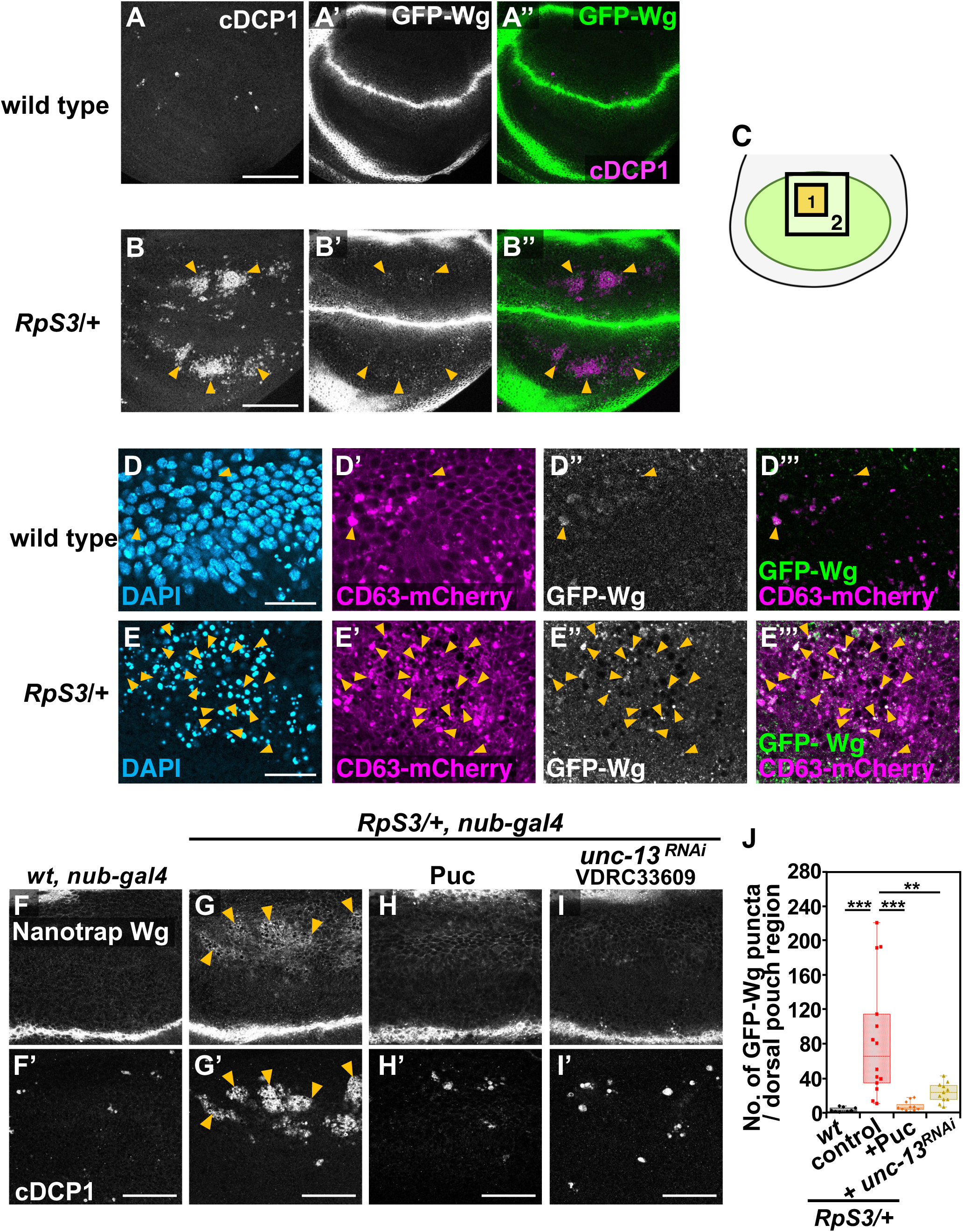
Dying cells secrete Wg via exocytosis in the *M/+* wing pouch. **(A-B’’)** Wing discs of *GFP-Wg/*+ (A), or *RpS3/+, GFP-Wg/+* (B) flies, and dying cells in the wing pouch were visualized by anti-cleaved Dcp-1 staining (white). GFP-Wingless was visualized by anti-GFP staining (white). Arrowheads indicate massive cell death in the *RpS3/+* wing pouch. Scale bar, 100 μm. **(D-E”’)** Immunofluorescent localization of the exosome marker CD63-mCherry, as visualized by anti-dsRed staining (magenta), and GFP-Wingless (white) in the wing pouch of *GFP-Wg/+* (D), or *RpS3/+*, *GFP-Wg/+* (E) flies expressing CD63-mCherry. The nuclei were visualized by DAPI staining (blue). The images correspond to the area enclosed by rectangle 1 in the schematic diagram of the wing disc (C). Orange arrowheads indicate the colocalization of GFP-Wingless-positive puncta with CD63-mCherry-positive puncta. Scale bar, 20μm. **(F-I’)** The membrane-tethered anti-GFP nanobody (Vhh4-CD8-HA) was expressed in the wing pouch of *GFP-Wg/+* (F), *RpS3/+*, *GFP-Wg/+* (G), *RpS3/+*, *GFP-Wg/+*, *nub-Gal4, UAS-Puc* (I), or *RpS3/+*, *GFP-Wg/+*, *nub-Gal4, UAS-unc-13-RNAi* (I) flies. GFP-Wingless was visualized by anti-GFP staining (white). Dying cells were visualized by anti-cleaved Dcp-1 staining in the wing discs (white). The images correspond to the area enclosed by the rectangle 2 in the schematic diagram of the wing disc (C). Immobilized Wg-GFP-positive puncta were frequently observed in the area of morphological dying cells in the *RpS3/+* wing pouch (Indicated by orange arrowheads). Scale bar, 50μm. **(J)** Boxplot with individual dots representing the number of GFP-Wingless-positive puncta per rectangle-2 in the wing pouch (C). The respective genotypes are shown in (F) (n=9), (G) (n=14), (H) (n=11), and (J) (n=11). Error bars, SEM; ***, p<0.001, **, p<0.01; Wilcoxon rank-sum test.

### JNK signaling generally triggers exocytosis-mediated Wg secretion

Finally, we investigated whether JNK activation generally upregulates exocytosis and subsequent Wg secretion in contexts beyond the *M/+* wing pouch. We generated JNK-activating clones by overexpressing Eiger, the *Drosophila* homolog of tumor necrosis factor (TNF), in the eye-antennal discs. Although dying cells were observed in Eiger-overexpressing clones (Figure 4G, compared to Figure 4F), the tissue size remained unchanged (data not shown), suggesting that neighboring cells proliferate to maintain tissue homeostasis. We found that the elimination of Eiger-expressing clones and the number of dying cells were significantly suppressed by knocking down *unc-13* in these clones (Figures 4D and 4H, quantified in Figure 4E and 4I), whereas unc-13-RNAi expression alone did not affect tissue growth (Figures 4A and 4C, quantified in Figure 4E). In addition, we observed a significant increase in Hrs-positive vesicles and GFP-Wg-positive puncta, derived from a knock-in allele ^34^, in Eiger-overexpressing clones compared to the control (Figure 4G’ and 4L’’, quantified in Figure 4J and 4M). Furthermore, the elimination of Eiger-expressing clones was significantly suppressed when the Wg signaling gradient was reduced either by deleting one copy of the *wg* gene (Figure 4N-4, quantified in Figure 4Q), suggesting the intriguing possibility that exocytosis-mediated Wg secretion from dying cells may be essential for both their cell death and proliferation of neighboring cells. Together, these data suggest that JNK signaling generally triggers exocytosis and the subsequent Wg secretion, which is crucial for maintaining tissue homeostasis.

**Figure 4.**
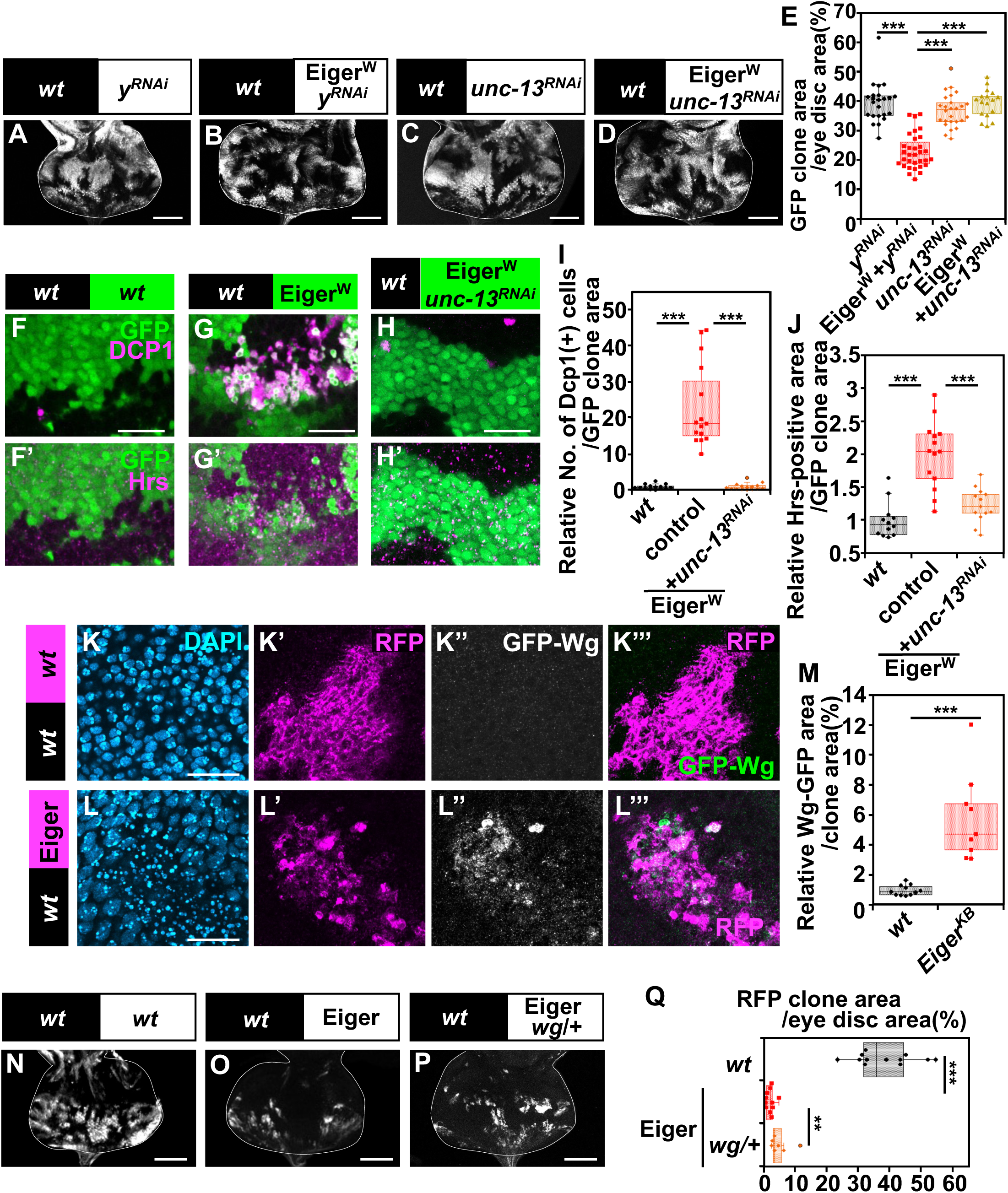
JNK signaling generally triggers exocytosis-mediated Wg secretion. **(A-D)** Eye disc bearing eyFLP-induced MARCM clones of *UAS-yellow-RNAi* (A), *UAS-Eiger^W^* +*UAS-yellow-RNAi* (B), *UAS-unc-13-RNAi* (C), or *UAS-Eiger^W^ +UAS-unc-13-RNAi* (D) cells. Scale bar, 100μm. **(E)** Boxplot with individual dots representing the clone size (% of total clone area per eye disc area) in genotypes shown in (A) (n=23), (B) (n=32), (C) (n=26), and (D) (n=17). Error bars, SEM; ***, p<0.001; Wilcoxon rank-sum test. **(F-H’)** Eye disc bearing eyFLP-induced MARCM clones of *wild-type* (F), *UAS-Eiger^W^* (G), or *UAS-Eiger^W^ +UAS-unc-13-RNAi* (H) cells stained with anti-cleaved Dcp-1 and anti-Hrs antibody (magenta). Scale bar, 20μm. **(I)** Boxplot with individual dots representing the number of cleaved-Dcp-1-positive dying cells per clone area in genotypes shown in (F) (n=14, number of clones), (G) (n=15), and (H) (n=15). Error bars, SEM; ***, p<0.001; Wilcoxon rank-sum test. **(J)** Boxplot with individual dots representing the relative Hrs-positive area in the eye disc for genotypes shown in (F’) (n=12, number of clones), (G’) (n=14), and (H’) (n=13). Error bars, SEM; ***, p<0.001; Wilcoxon rank-sum test. **(K-L’’’)** Eye disc bearing eyFLP-induced MARCM clones of *GFP-Wg/*+ (K), or *UAS-Eiger^W^*, *GFP-Wg/*+ (L) cells stained with anti-GFP antibody (white). Scale bar, 20μm. **(M)** Boxplot with individual dots representing the relative GFP-Wg area per clone area of the eye disc for genotypes shown in (K) (n=11), and (L) (n=9). Error bars, SEM; ***, p<0.001; Wilcoxon rank-sum test. **(N-P)** Eye disc bearing eyFLP-induced MARCM clones of wild-type (N), *UAS-Eiger^KB^* (O), or *UAS-Eiger^KB^*, *wg/+* (P) cells. Scale bar, 100μm. **(Q)** Boxplot with individual dots representing the clone size (% of total clone area per eye disc area) in genotypes shown in (N) (n=12), (O) (n=11), and (P) (n=7). Error bars, SEM; ***, p<0.001, **, p<0.01; Wilcoxon rank-sum test.

## Discussion

Our current study reveals that exocytosis is crucial for massive cell-turnover in *M/+* wing morphogenesis, acting as a downstream event of JNK signaling. In dying cells, JNK signaling upregulates the expression of exocytosis-related genes and Wg, leading to increased Wg secretion. In addition, elevated exocytosis induces cell death via an exocytosis-Dronc amplification loop (Figure 5). Notably, impairment of exocytosis in dying cells in *RpS3/+* animals leads to phenotypic variations in adult wings (Figures 2O and 2P in this study), suggesting that increased exocytosis in dying cells, followed by subsequent cell-turnover, may provide the flexibility needed to adjust developmental programs.

**Figure 5.**
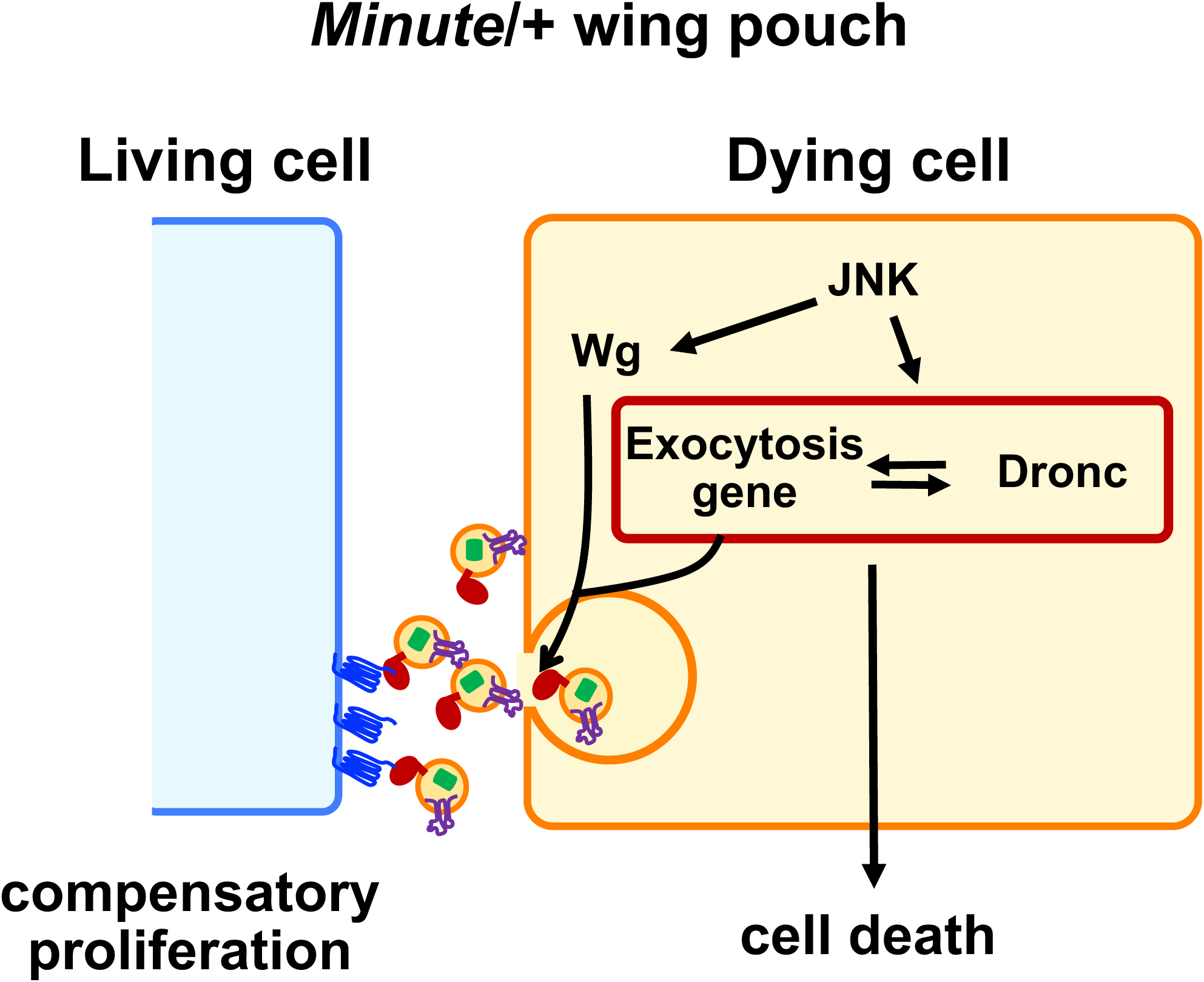
A model for ensuring robust coordination of tissue growth in *M*/+ animals. During the larval period of *M*/+ animals, JNK signaling is activated in the *M/+* wing pouch, which induces the expression of exocytosis-related genes and Wg, leading to the secretion of Wg through exocytosis in dying cells. In addition, elevated exocytosis induces cell death via an exocytosis-Dronc amplification loop. This process subsequently promotes the proliferation of neighboring cells, establishing their apoptotic/proliferative status and resulting in significant cell turnover in the *M*/+ wing pouch.

We identified Wg as an important factor secreted through exocytosis from dying cells, potentially essential for the proliferation of neighboring living cells to maintain tissue homeostasis. Previous studies have shown that ‘undead’ cells, stimulated toward apoptosis but kept alive by the caspase inhibitor p35, are able to promote the proliferation of neighboring cells in the *Drosophila* epithelium ^57–59^. Several mitogens including Wg, Dpp, and Spitz (a EGF homolog) have been identified as being produced by apoptotic cells in a JNK-dependent manner ^18,51,60^. In addition, Hedgehog (Hh) ^15^ have also been identified as products of apoptotic cells. Furthermore, during cell competition between wild-type (winners) and *M/+* cells (losers) in the *Drosophila* epithelium, prospective loser cells secrete upd3, which promotes competitive elimination by winner cells ^12^. Hence, in addition to Wg, other mitogens could also be secreted thorough exocytosis from dying cells, potentially contributing to massive cell-turnover in the *M/+* wing pouch.

Previous studies have identified exosomes as carriers of Wnt/Wg in the extracellular space of both mammalian and *Drosophila* cells, including wing disc cells ^28,53–56,61^. Concurrently, alternative mechanisms for Wg transport, such as those involving lipoprotein particles or a lipocalin Swim in the *Drosophila* wing disc, have been reported ^29,62^. Additionally, the cell-surface proteoglycan Dally-like-protein (Dlp) has been reported to enable long-range signaling of the palmitoylated Wg ^34^. Intriguingly, Wnt/Wg transport has also been observed through filopodia-like cellular extensions known as cytonemes ^63–66^. In our study of the *M*/+ wing disc, a model characterized by massive cell-turnover, we observed partial colocalization of Wg-GFP puncta with exosomal markers such as CD63-mCherry and Hrs. Our data also suggest that Wg secretion through exocytosis may not uniformly occur among all dying cells within this context. This disparity suggests that while exosomes from dying cells significantly contribute to Wg transport within the *M*/+ wing pouch, other pathways may also be operative.

*Rp* genes are crucial in a broad range of organisms, from yeast to humans, frequently exhibiting haploinsufficient mutant phenotypes. Notably, heterozygous mutations in genes encoding ribosomal proteins or ribosomal biogenesis factors are linked to tissue-specific human diseases known as ribosomopathies ^67^. In patients affected by these conditions, some tissues may preserve normal patterning and functions, potentially by buffering stresses caused by ribosomal mutations. Given the evolutionary conservation of the molecules we identified in *Drosophila*, regulating cell turnover through exocytosis in dying cells may represent evolutionarily conserved strategy to prevent developmental abnormalities in ribosomal protein mutants.

## Author Contributions

**Conceptualization:** Nanami Akai, Tatsushi Igaki, Shizue Ohsawa.

**Data curation:** Nanami Akai, Tatsushi Igaki, Shizue Ohsawa.

**Formal analysis:** Nanami Akai.

**Funding acquisition:** Shizue Ohsawa, Tatsushi Igaki.

**Investigation:** Nanami Akai, Shizue Ohsawa.

**Methodology:** Nanami Akai.

**Project administration:** Shizue Ohsawa.

**Resources:** Shizue Ohsawa.

**Supervision:** Shizue Ohsawa.

**Visualization:** Nanami Akai, Tatsushi Igaki, Shizue Ohsawa.

**Writing – original draft:** Shizue Ohsawa, Nanami Akai.

**Writing – review & editing:** Nanami Akai, Tatsushi Igaki, Shizue Ohsawa.

## Acknowledgments

We thank Takefumi Kondo, Keisuke Ikawa, Kiichiro Taniguchi, Rina Nagata, and Yukari Sando for discussions, Susumu Tsutsumi, Nana Watanabe, Mina Hoshino, Tomoko Furukawa, and Sayako Suzuki for technical support, and NGS core facility of the Graduate Schools of Biostudies, Kyoto University for supporting the RNA-seq analysis. We also thank Yash Hiromi, Gary Struhl, Maoto Sato, Jean-Paul Vincent, Sharad Kumar, Konrad Basler, the Bloomington *Drosophila* Stock Center (Indiana), the Vienna *Drosophila* Resource Center (Vienna), the National Institute of Genetics Stock Center (Mishima), and the *Drosophila* Genomics and Genetic Resources (Kyoto) for fly stocks. Cell sorting using BD FACS Aria II were performed at the Medical Research Support Center, Graduate School of Medicine, Kyoto University, supported by Basis for Supporting Innovative Drug Discovery and Life Science Research (BINDS) from AMED (Grant No. JP22ama121034). This work was supported in part by MEXT Grant-in-Aid (KAKENHI) for Transformative Research Area (A) (Grant Nos. 20H05945 to SO, 21H05284 to TI), the Scientific Research (B) (Grant No. 22H02616 to SO), and Challenging Exploratory Research (Grant No. 21K19257 to SO), Japan Science and Technology Agency (Moonshot Research & Development: Grant Number JPMJPS2022 to SO), AMED-CREST, Japan Agency for Medical Research and Development (22gm1710002h0001 to TI), and Japan Agency for Medical Research and Development (21gm5010001 to TI) .

## STAR Methods

### Resource availability

#### Lead contact

Further information and requests for resources and reagents should be directed to and will be fulfilled by the lead contact, Shizue Ohsawa.

#### Materials availability

*Drosophila* lines generated in this study are available from the lead contact without restriction.

### Experimental model and subject detail

#### Fly strains

Fly stocks were cultured at 25°C on standard fly food. To detect dying cells in the wing pouch, the following strains were used: *nub-Gal4; UAS-CD8-PARP-Venus* (control); *nub-Gal4; UAS-CD8-PARP-Venus, RpS3^Plac^*^92^ (BL5627)*/TM6B* (*RpS3*/+ tester). Additional strains were used as follows: *UAS-EGFP.CD63* (BL91390), *UAS-CD63.mCherry* (BL91389), *UAS-Syt1-eGFP* (BL6925), *RpL19^K^*^03704^ (DGRC102285), *20XUAS-IVS-GCaMP6m* (BL42750), *UAS-unc-13-RNAi* (VDRC33609, NIG2999R-2), *UAS-LexA-RNAi* (BL67947), *UAS-Rab3-RNAi* (NIG7576R-3, BL31691), *UAS-SNAP25-RNAi* (HMS01367, BL27306), *UAS-Syt1-RNAi* (NIG3139R-1, BL31668), *UAS-Rab27-RNAi* (NIG14791R-2, BL31887), *UAS-ALiX-RNAi* (HMS00298, VDRC32049), *Gαq*^221^*^C^* (BL30744), *UAS-Gαq-RNAi* (NIG17759R-2), *UAS-Plc21C-RNAi* (NIG4574R-2, 4574R-3), *norpA*^7^ (DGRC108362)*, UAS-norpA-RNAi* (VDRC105676), *UAS-y-RNAi* (NIG3757R-1), *wg^l-^*^8^ (BL5351), *UAS-Puc* ^68^, *UAS-Dronc^DN^*(Sharad Kumar) ^69^*, GFP-Wingless^S^*^239^*^A^* (JP. Vincent) ^34^, *UAS-VHH4-CD8-HA* (JP. Vincent) ^34^, *UAS-Eiger^+W^* ^70^, and *UAS-Eiger^KB^* (Konrad Basler) ^71^.

### Method details

#### Histology

Wandering third instar larvae were dissected in phosphate buffered saline (PBS) and fixed with 4%-Paraformaldehyde (PFA) for 5 min on ice, followed by 20 min at room temperature. The fixed larvae were washed 3 times with PBT (PBS + 0.1% TritonX100) for 20 min each, and were then blocked with PBTn (5% donkey serum + PBT) solution for 30 min. For immunostaining, they were incubated overnight at 4°C with primary antibodies in PBTn. After four 30-min washes with PBT, the samples were re-blocked in PBTn for 30 minutes, then incubated with secondary antibodies for 2 hours at room temperature. Following another set of four 30-minute washes with PBT, the samples were mounted using either SlowFade Diamond Antifade Mountant (Invitrogen, #S36972) or an alternative anti-fade mounting medium. The latter was composed of 70% Glycerol (Sigma, #G5516), 0.2% n-Propyl Gallate (KANTO CHEMICAL, #32465-31), and supplemented with DAPI (Sigma, #D9542). For extracellular immunostaining, larvae were dissected in ice-cold Schneider’s *Drosophila* Medium (Gibco, #21720024) and incubated with primary antibodies for 2 hours on ice. The larvae were then rinsed with ice-cold Schneider’s *Drosophila* Medium followed by PBS and fixed with 4% PFA for 40 min at room temperature. After fixation, samples were washed with PBS. Subsequent processing was the same as in the preceding protocol. Images were taken with Leica SP5, Leica SP8, Leica SPE, Leica STELLARIS and ZEISS LSM900 confocal microscopes. Primary antibodies used are as follows; rat anti-GFP antibody (Nacalai Tesque, #04404-26, 1:1000) (Extracellular GFP, 1:300), rabbit anti-cleaved PARP (Asp214) antibody (Cell Signaling Technology (CST), #9541, 1:200), rabbit anti-phospho-histone H3 (Ser10) antibody (CST, #9706, 1:100), rabbit anti-DsRed antibody (Takara Bio Clontech (CLN), #632496, 1:250) (Extracellular DsRed, 1:75), mouse anti-Hrs antibody (DSHB, #27-4-s, 1:250), and rabbit anti-Cleaved *Drosophila* Dcp-1 (Asp216) antibody (CST, #9678, 1:100) for 3 days. Secondary antibodies used are as follows; Goat anti-rabbit Alexa 488 (1:250, Invitrogen #A11034), Goat anti-rabbit Alexa 546 (1:250, Invitrogen #A11035), Goat anti-rat Alexa 488 (1:250, Invitrogen #A11006), Goat anti-chicken Alexa 488 (1:250, Invitrogen, #A11039), and Goat anti-mouse Alexa 647 (1:250, Invitrogen #A21237) antibodies.

### Quantification of puncta number

The vesicle marker EGFP-CD63, CD63-mCherry, and Syt1-eGFP were expressed in the wing pouch under the control of *nub-gal4 driver*. In the region where massive cell death is observed in the *M/+* wing disc, these markers, in addition to GFP-Wingless signals, were acquired at the confocal plane exhibiting the highest signal intensity.

CD63-mCherry were also expressed within the mosaic clones of the eye-antenna disc. Puncta were manually counted using Fiji software and the data were analyzed with KaleidaGraph (Synergy Software).

### Quantitative Analysis of GFP-Wingless and CD63-mCherry Colocalization

To analyze the overlap between GFP-Wingless-positive puncta and CD63-mcherry-positive vesicles, their colocalization was manually quantified in the regions exhibiting massive cell death. This quantification was conducted using the confocal plane that displayed the highest signal intensity. Images were manually thresholded to eliminate background signals, and the areas of overlap were calculated using Fiji software, followed by further analyses with KaleidaGraph.

### Calcium Time-lapse imaging and quantification

To monitor Ca^2+^ activity, wandering third instar larvae were dissected and cultured in Schneider’s *Drosophila* medium supplemented with 2% FBS, 1% Insulin solution (Sigma-Aldrich, #I0516), 3% fly extract (VDRC, #VFE1.25), and 2% low-melting agarose (Sigma-Aldrich, #A4018)/PBS. Imaging was conducted at 10 sec intervals for 20-min with a Leica STELLARIS confocal microscope ^42,72^. For quantitative analysis, dying cells exhibiting continuous calcium activity were manually counted using Fiji software and the data were analyzed with KaleidaGraph.

### Detection of dying cells and quantification using CD8-PARP-Venus

To detect dying cells, the activated-caspase-3 indicator CD8-PARP-Venus was expressed in the pouch under the control of *nub-gal4* driver. Cleaved PARP signals were acquired at the confocal plane where signal intensity was highest. The number of these signals in the wing pouch was manually measured using Fiji software and the data were analyzed with KaleidaGraph.

### Quantification of pH3-positive area

For the analysis of pH3-positive cells, total area of pH3-positive cells and the size of the wing pouch area were automatically measured from confocal slices of Z-stack images compressed using maximum projection functions in Fiji software, as previously described ^73^. The acquired data were then analyzed with KaleidaGraph.

### Measurement of adult wings

Left and right wings of female flies were rinsed with xylene and mounted in a Canada Balsam (Nacalai Tesque). Images of the wings were acquired using Leica M205C stereo microscope.

### Quantification of clone size

For the analysis of clone size in the eye disc, total clone area as a percentage of the disc area (%) was measured from XY confocal images using Fiji software, as previously described ^73^. The acquired data were then analyzed with KaleidaGraph.

### FACS and mRNA-seq analysis

GFP+ wing pouch cells were isolated using a FACS Aria II cell sorter (BD Bioscience). Total RNA was then extracted for RNA sequencing analysis, following the method previously described ^74^. Briefly, third instar larvae were dissected in ice-cold PBS. The wing disc cells were dissociated using Trypsin (10x TrypLE Select) (Gibco, #A1217701) at 37°C for 20 min. The enzymatic reaction was stopped with Schneider’s medium containing 5% FBS, and GFP-positive cells from wing pouches were sorted using a FACS Aria II cell sorter (BD Bioscience). RNA samples were extracted using NucleoSpin RNA XS Kit (TaKaRa, #740902). The quality of the extracted total RNA was assessed using an Agilent 2100 Bioanalyzer, and then Strand-specific Libraries for mRNA-seq were prepared using the NEBNext Ultra II Directional RNA Library Prep Kit for Illumina (E7760S). For RNA-seq, NextSeq 500/550 High output Kit v2.5 was utilized on a NextSeq 500 (Illumine), generating single-end reads at a length of 85 bases.

### RNA-seq data analysis

Each genotype was analyzed across three independent replicates. Each sequencing experiment generated more than 16 million raw reads. These reads were then quality-filtered using Trim_Galore! (v0.6.5) (https://www.bioinformatics.babraham.ac.uk/projects/ trim_galore/) with the default setting to remove low-quality reads and adaptor sequences. The filtered reads were mapped to the *Drosophila* melanogaster genome (Ensembl BDGP6.32), obtained from Illumina iGenomes using STAR ^75^. In the *RpS3/+, nub-Gal4, UAS-Puc* experiment, one sample exhibited only 65.69% of reads being uniquely mapped, which was lower compared to other experiments where more than 80% of reads were uniquely mapped in each sample. The number of reads mapped to each gene was quantified using RSEM (v1.3.3) ^76^. Normalization was performed with calcNormFactors function in edgeR (v3.30.3) ^77^. Significantly differentially expressed genes were identified with an FDR<0.05 using the glmQLFit and glmQLFTest function in edgeR (R version 4.0.1, limma 3.44.3). Furthermore, GO enrichment analysis was performed on the 362 genes regulated by JNK signaling, utilizing the online tool available at “Gene Ontology” (https://geneontology.org).

### Statistical analysis

Statistical analysis was performed, and boxplot graphs were generated using KaleidaGraph ver4.5 (Synergy Software). All experiments represented in the same graph were conducted at the same time. In bar graphs, error bars indicate the standard deviation (SD). Significance levels are indicated as follows: * P<0.05, ** P<0.01, and *** P<0.001. All statistical data were summarized in Table S4.

## Supplemental information

**Document S1. Figures S1-S3 and Table S5.**

**Video S1. Calcium Time-lapse imaging of wing discs. Related to Figure 1**.

The calcium reporter GCaMP6m was expressed in wing pouches of wild-type larvae. Wing discs were dissected and subjected to time-lapse imaging at 10-second intervals with a Leica STELLARIS confocal microscope. See *Methods* for details.

**Video S2. Calcium Time-lapse imaging of wing discs. Related to Figure 1**.

The calcium reporter GCaMP6m was expressed in wing pouches of *RpS3/+* larvae. Wing discs were dissected and subjected to time-lapse imaging at 10-second intervals with a Leica STELLARIS confocal microscope. See *Methods* for details.

**Video S3. Calcium Time-lapse imaging of wing discs. Related to Figure 1**.

The calcium reporter GCaMP6m was expressed in wing pouches of *RpS3/+*, *nub-Gal4, UAS-Puc* larvae. Wing discs were dissected and subjected to time-lapse imaging at 10-second intervals with a Leica STELLARIS confocal microscope. See *Methods* for details.

**Table S1 List of genes identified as differentially expressed by RNA-seq between RpS3/+ and WT, and between *RpS3*/+, +Puc and WT pouch cells.**

**Table S2 List of genes differentially expressed in *RpS3*/+ pouch cells, associated with JNK or JAK/STAT targets, Gr64 gustatory receptors, oxidative stress, and DNA repair.**

**Table S3 List of genes identified as differentially expressed by RNA-seq between *RpS3*/+ and *RpS3*/+, +Puc pouch cells.**

**Table S4 Summary of statistical analysis.**

