## Supplementary material for "JNK signaling-mediated exocytosis coordinates epithelial cell-turnover in *Drosophila* ribosomal protein mutants": Figures S1-S3 and Table S5

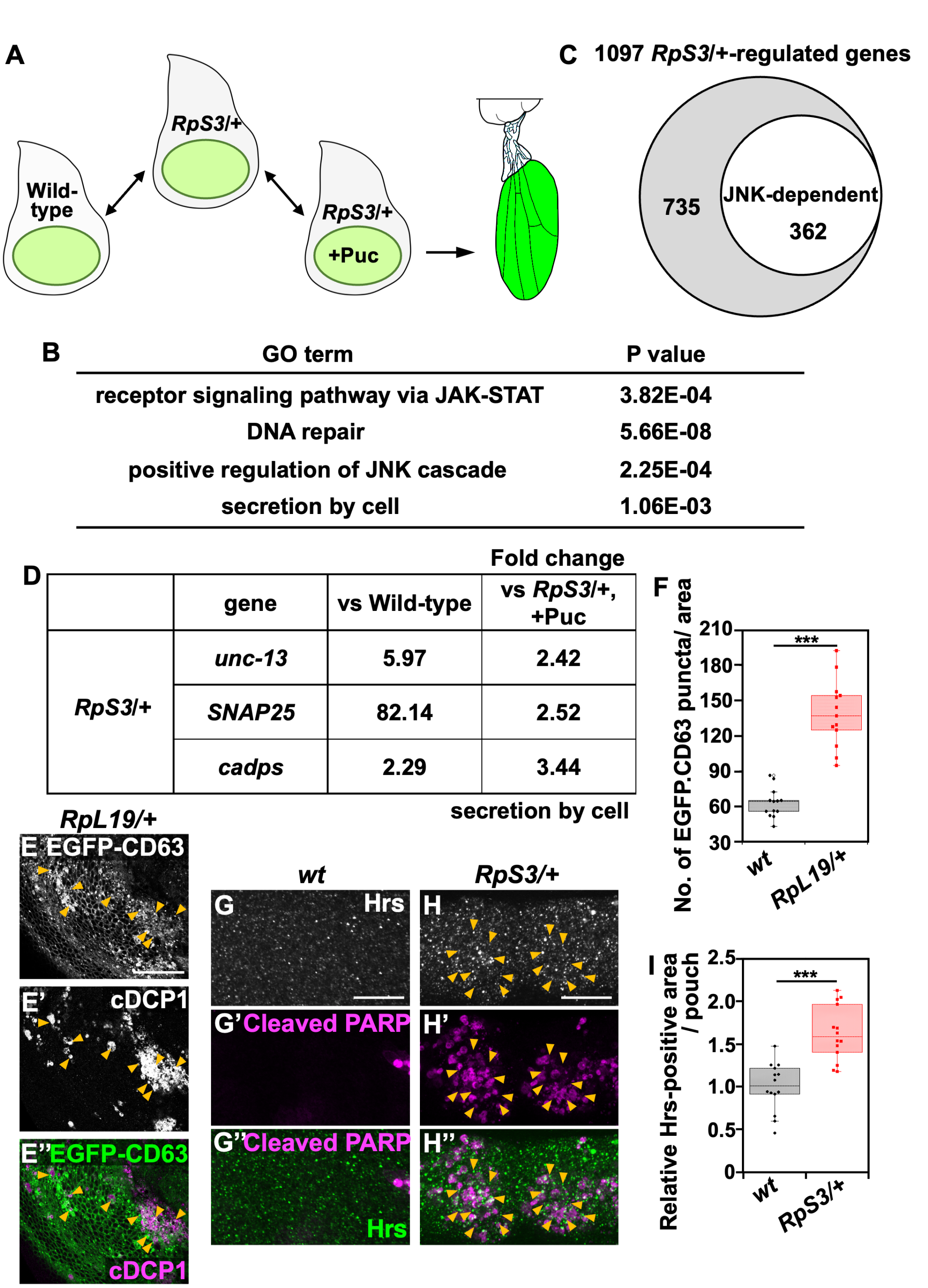

**Figure S1.** ***M/+*** **wing pouch elevates exocytosis downstream of JNK signaling. Related to Figure 1.**

**(A)** Schematic representation of the genotypes used for transcriptional profiling: mRNA was isolated from FACS-Sorted GFP$^{+}$ pouch cells (green) in third instar wing discs of wild-type, *RpS3*/+, or *RpS3*/+, *nub-Gal4, UAS-Puc* flies.

**(B)** Four GO terms for biological process were significantly enriched in the *RpS3/*+-regulated genes.

**(C)** Venn diagram depicting genes that are differentially expressed in *RpS3*/+ cells compared to wild-type cells, with 362 of these changes being regulated by JNK signaling.

**(D)** Expression levels (fold changes relative to wild-type control or *RpS3*/+, *nub-Gal4, UAS-Puc*) for *RpS3*/+ -regulated genes associated with the "secretion by cell" GO term. These genes were regulated by JNK signaling in the *RpS3/*+ wing pouch.

**(E)** EGFP-CD63 was expressed in the wing pouch of *RpL19/*+ flies (white). Dying cells were visualized by anti-cleaved Dcp-1 staining in the wing discs (white). Arrowheads indicate massive cell death in the *RpL19/*+ wing pouch. Scale bar, 50 μm.

**(F)** Boxplot with individual dots representing the number of EGFP-CD63-positive puncta in the pouch of genotypes shown in (Figure 1A) (n=13, number of wing pouches), and (E) (n=13). Error bars, SEM; ***, p<0.001; Wilcoxon rank-sum test.

**(G-H'')** The activated-caspase-3 indicator CD8-PARP-Venus was expressed in the wing pouch of wild-type (G), or *RpS3/*+ (H) flies, and dying cells in the wing pouch were visualized by anti-cleaved PARP staining (magenta). Hrs expression was visualized by anti-Hrs staining (white). Hrs-positive vesicles were frequently observed in the area of morphological dying cells in the *RpS3/*+ wing pouch (Indicated by orange arrowheads). The images correspond to the area enclosed by rectangle 1 in the schematic diagram of the wing disc (Figure 3C). Scale bar, 20 μm.

**(I)** Boxplot with individual dots representing the number of Hrs-positive puncta in the pouch of genotypes shown in (G) (n=14, number of wing pouches), and (H) (n=14). Error bars, SEM; ***, p<0.001; Wilcoxon rank-sum test.

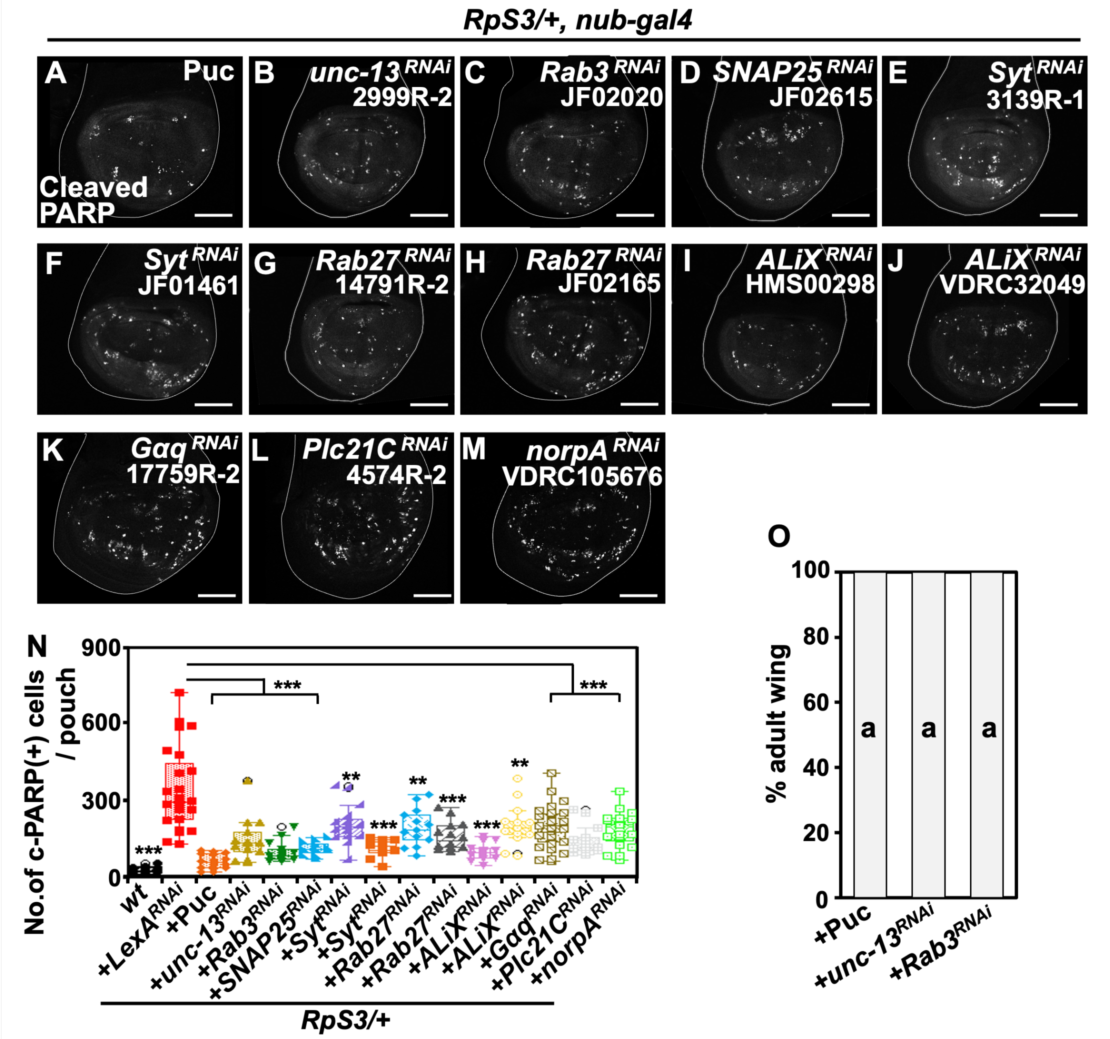

**Figure S2. Exocytosis is required for massive cell-turnover in the *M/+* wing pouch. Related to Figure 2.**

**(A-M)** CD8-PARP-Venus was expressed in the wing pouch of *RpS3/+, nub-Gal4, UAS-Puc* (A), *RpS3/+, nub-Gal4, UAS-unc-13-RNAi* (B), *RpS3/+, nub-Gal4, UAS-Rab3-RNAi* (C), *RpS3/+, nub-Gal4, UAS-SNAP25-RNAi* (D), *RpS3/+, nub-Gal4, UAS-Syt-RNAi* (E and F), *RpS3/+, nub-Gal4, UAS-Rab27-RNAi* (G and H), *RpS3/+, nub-Gal4, UAS-ALiX-RNAi* (I and J), *nub-Gal4, UAS-Gaq-RNAi* (K), *nub-Gal4, UAS-Plc21C-RNAi* (L), or *RpS3/+, nub-Gal4, UAS-norpA-RNAi* (M) flies, and dying cells in the wing pouch were visualized by anti-cleaved PARP staining (white). Scale bar, 100μm.

**(N)** Boxplot with individual dots representing the number of cleaved-PARP-positive dying cells per pouch in genotypes shown in (Figure 2F) (n=18), (2G) (n=24), (A) (n=18), (B) (n=14), (C) (n=13), (D) (n=12), (E) (n=15), (F) (n=10), (G) (n=13), (H) (n=14), (I) (n=13), (J) (n=13), (K) (n=29), (L) (n=14), and (M) (n=19). Error bars, SEM; ***, p<0.001, **, p<0.01; Wilcoxon rank-sum test.

**(O)** The rate of defective wings in the genotypes of *nub-Gal4, UAS-Puc* (n=41), *nub-Gal4, UAS-unc-13-RNAi* (n=55), and *nub-Gal4, UAS-Rab3-RNAi* (n=61).

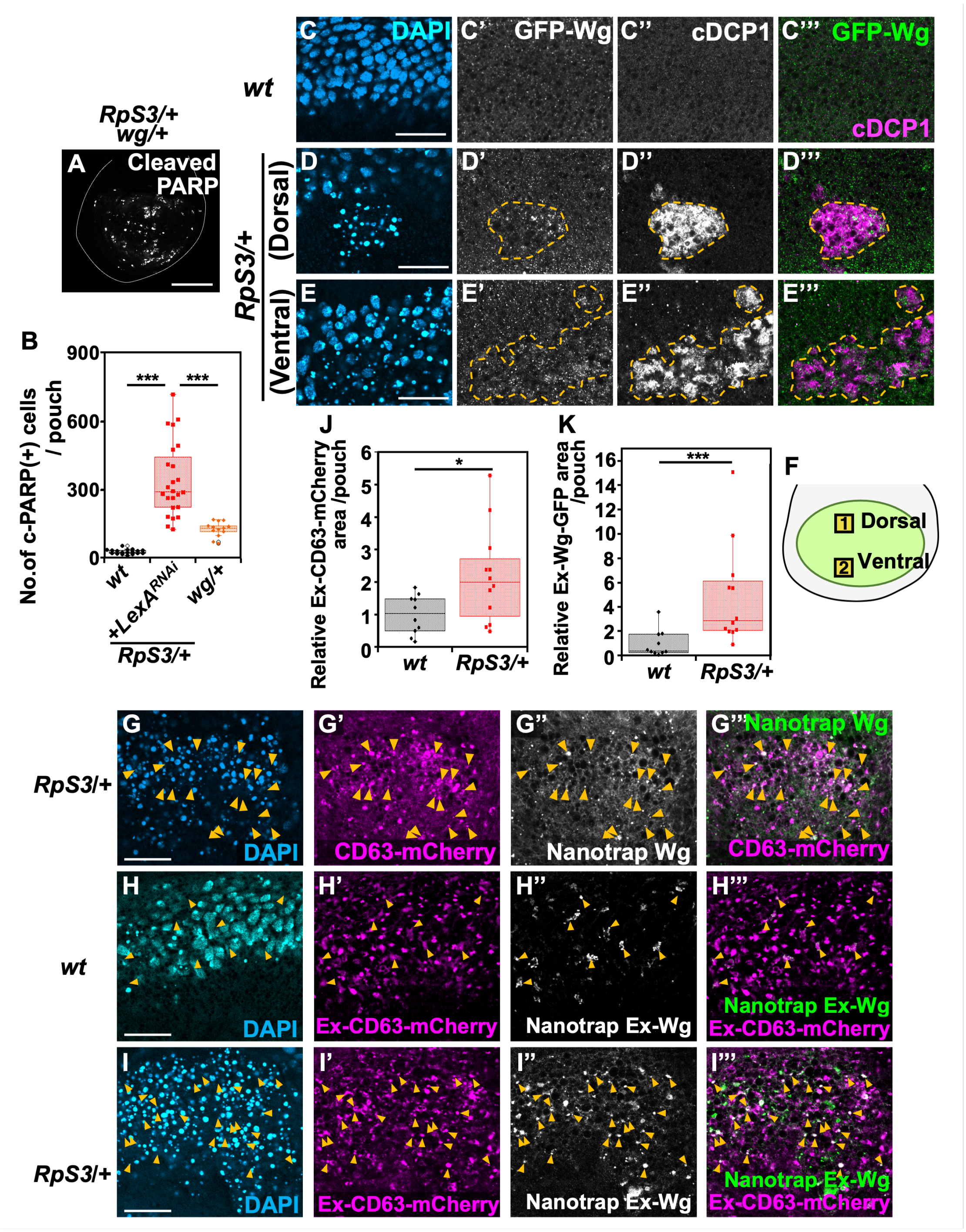

**Figure S3. Dying cells Secrete Wg via exocytosis in the *M*/+ wing pouch. Related to Figure 3.**

**(A)** The activated-caspase-3 indicator CD8-PARP-Venus was expressed in the wing pouch of *RpS3/+, wg/+* flies, and dying cells in the wing pouch were visualized by anti-cleaved PARP staining (white). Scale bar, 100μm.

**(B)** Boxplot with individual dots representing the number of cleaved-PARP-positive dying cells per pouch in genotypes shown in (Figure 2H) (n=18), and (A) (n=13). Error bars, SEM; ***, p<0.001; Wilcoxon rank-sum test.

**(C-E''')** Wing discs from *GFP-Wg*/+ (C), or from the dorsal (D) or ventral (E) regions of the wing disc in *RpS3*/+, *GFP-Wg*/+ flies. The images correspond to the areas enclosed by rectangle 1 and rectangle 2, respectively, in the schematic diagram of the wing disc (F). Dying cells in the wing pouch were stained with anti-cleaved Dcp-1 antibody (white). GFP-Wingless was visualized by anti-GFP staining (white). The regions outlined by dashed lines indicate massive cell death in the *RpS3/*+ wing pouch. Scale bar, 20μm.

**(G)** CD63-mcherry and Vhh4-CD8-HA were expressed in the wing pouch of *RpS3/*+, *GFP-Wg/*+ flies. CD63-mCherry was visualized by anti-dsRed staining (magenta). GFP-Wingless was visualized by anti-GFP staining (white). The nuclei were visualized by DAPI staining (blue). Orange arrowheads indicate the colocalization between CD63-mCherry-positive and GFP-Wingless-positive puncta. The images correspond to the area enclosed by rectangle 1 in the schematic diagram of the wing disc (Figure 3C). Scale bar, 20μm.

**(H-I''')** Vhh4-CD8-HA was expressed in the wing pouch of *GFP-Wg/*+ (H), or *RpS3/*+, *GFP-Wg/*+ (I) flies. Extracellular CD63-mCherry and extracellular GFP-Wingless were visualized by anti-dsRed staining (magenta) and anti-GFP staining (white), respectively. The nuclei were visualized by DAPI staining (blue). Orange arrowheads indicate the colocalization of extracellular GFP-Wingless-positive puncta with extracellular CD63-mCherry-positive puncta. The images correspond to the area enclosed by the rectangle 1 in the schematic diagram of the wing disc (Figure 3C). Scale bar, 20μm.

**(J)** Boxplot with individual dots representing extracellular CD63-mCherry-positive area in the region enclosed by rectangle 1 in the schematic diagram of the wing disc (Figure 3C). The respective genotypes are shown in (H) (n=10), and (I) (n=12). Error bars, SEM; *, p<0.05; Wilcoxon rank-sum test.

**(K)** Boxplot with individual dots representing extracellular GFP-Wingess-positive area in the region enclosed by rectangle 1 in the schematic diagram of the wing disc (Figure 3C). The respective genotypes are shown in (H) (n=10), and (I) (n=12) Error bars, SEM; ***, p<0.001; Wilcoxon rank-sum test.

| Figure 1 | |
|---|---|
| A | *nub-Gal4/UAS-EGFP.CD63; +/+* |
| B | *nub-Gal4/UAS-EGFP.CD63; RpS3$^{Plac92}$/+* |
| C | *nub-Gal4/UAS-EGFP.CD63; RpS3$^{Plac92}$/UAS-Puc* |
| D | *nub-Gal4/UAS-EGFP.CD63; RpS3$^{Plac92}$/UAS-unc-13-RNAi (VDRC33609)* |
| G | *nub-Gal4/UAS-Syt1-eGFP; +/+* |
| H | *nub-Gal4/ UAS-Syt1-eGFP; RpS3$^{Plac92}$/+* |
| I | *nub-Gal4/ UAS-Syt1-eGFP; RpS3$^{Plac92}$/UAS-Puc* |
| J | *nub-Gal4/ UAS-Syt1-eGFP; RpS3$^{Plac92}$/UAS-unc-13-RNAi (VDRC33609)* |
| L | *nub-Gal4/+; 20XUAS-IVS-GCaMP6m/+* |
| M | *nub-Gal4/+; 20XUAS-IVS-GCaMP6m/RpS3$^{Plac92}$* |
| N | *nub-Gal4/+; 20XUAS-IVS-GCaMP6m/RpS3$^{Plac92}$, UAS-Puc* |

| Figure 2 | |
|---|---|
| A | *nub-Gal4/+; UAS-CD8-PARP-Venus/+* |
| B | *nub-Gal4/+; UAS-CD8-PARP-Venus, RpS3$^{Plac92}$/+* |
| C | *nub-Gal4/+; UAS-CD8-PARP-Venus, RpS3$^{Plac92}$/UAS-Puc* |
| D | *nub-Gal4/+; UAS-CD8-PARP-Venus, RpS3$^{Plac92}$/UAS-unc-13-RNAi (VDRC33609)* |
| E | *nub-Gal4/UAS-Rab3-RNAi (7576R-3); UAS-CD8-PARP-Venus, RpS3$^{Plac92}$/+* |
| G | *nub-Gal4/+; UAS-CD8-PARP-Venus/+* |
| H | *nub-Gal4/+; UAS-CD8-PARP-Venus, RpS3$^{Plac92}$/UAS-LexA-RNAi (HMS05772)* |
| I | *nub-Gal4/+; UAS-CD8-PARP-Venus, RpS3$^{Plac92}$/UAS-unc-13-RNAi (VDRC33609)* |
| J | *nub-Gal4/UAS-Rab3-RNAi (7576R-3); UAS-CD8-PARP-Venus, RpS3$^{Plac92}$/+* |
| K | *nub-Gal4/+; UAS-CD8-PARP-Venus, RpS3$^{Plac92}$/UAS-SNAP25-RNAi (HMS01367)* |
| L | *nub-Gal4/ Gαq$^{221c}$ (BL30744); UAS-CD8-PARP-Venus, RpS3$^{Plac92}$/+* |
| M | *nub-Gal4/+; UAS-CD8-PARP-Venus, RpS3$^{Plac92}$/ UAS-Plc21C-RNAi (4574R-3)* |
| N | *norpA$^{7}$ (DGRC108362)/+ ; nub-Gal4/+; UAS-CD8-PARP-Venus, RpS3$^{Plac92}$/+* |
| O | |
| wt | *nub-Gal4/+; UAS-CD8-PARP-Venus/+* |
| ***RpS3*/+** | *nub-Gal4/+; UAS-CD8-PARP-Venus, RpS3$^{Plac92}$/+* |
| ***RpS3*/+, +Puc** | *nub-Gal4/+; UAS-CD8-PARP-Venus, RpS3$^{Plac92}$/UAS-Puc* |
| ***RpS3*/+, +unc-13-RNAi** | *nub-Gal4/+; UAS-CD8-PARP-Venus, RpS3$^{Plac92}$/UAS-unc-13-RNAi (VDRC33609)* |
| ***RpS3*/+, +Rab3-RNAi** | *nub-Gal4/UAS-Rab3-RNAi (7576R-3); UAS-CD8-PARP-Venus, RpS3$^{Plac92}$/+* |
| Q | *nub-Gal4/+; UAS-CD63.mCherry /+* |
| R | *nub-Gal4/+; UAS-CD63.mCherry, RpS3$^{Plac92}$/+* |
| S | *nub-Gal4/+; UAS-CD63.mCherry, RpS3$^{Plac92}$/UAS-Dronc$^{DN}$* |

| Figure 3 | |
|---|---|
| A | *GFP-Wingless /+* |
| B | *GFP-Wingless /+; RpS3$^{Plac92}$/+* |
| D | *nub-Gal4/GFP-Wingless; UAS-CD63.mCherry/+* |
| E | *nub-Gal4/GFP-Wingless; UAS-CD63.mCherry/RpS3$^{Plac92}$* |
| F | *nub-Gal4, UAS-VHH4-CD8-HA/GFP-Wingless* |
| G | *nub-Gal4, UAS-VHH4-CD8-HA/GFP-Wingless; RpS3$^{Plac92}$/+* |
| H | *nub-Gal4, UAS-VHH4-CD8-HA/GFP-Wingless; RpS3$^{Plac92}$/UAS-Puc* |
| I | *nub-Gal4, UAS-VHH4-CD8-HA/GFP-Wingless; RpS3$^{Plac92}$/UAS-unc-13-RNAi (VDRC33609)* |

| | |
|---|---|
| **Figure 4** | |
| **A** | *eyFLP1/+; TubGal80, FRT40A/FRT40A; Act>y[+]>Gal4, UAS-GFP/UAS-y-RNAi (3757R-1)* |
| **B** | *eyFLP1/+; TubGal80, FRT40A/FRT40A, UAS-Eiger*$^{+W}$*; Act>y[+]>Gal4, UAS-GFP/UAS-y-RNAi (3757R-1)* |
| **C** | *eyFLP1/+; TubGal80, FRT40A/FRT40A; Act>y[+]>Gal4, UAS-GFP/UAS-unc-13-RNAi (VDRC33609)* |
| **D** | *eyFLP1/+; TubGal80, FRT40A/FRT40A, UAS-Eiger*$^{+W}$*; Act>y[+]>Gal4, UAS-GFP/ UAS-unc-13-RNAi (VDRC33609)* |
| **F** | *eyFLP1/+; TubGal80, FRT40A/FRT40A; Act>y[+]>Gal4, UAS-GFP/+* |
| **G** | *eyFLP1/+; TubGal80, FRT40A/FRT40A, UAS-Eiger*$^{+W}$*; Act>y[+]>Gal4, UAS-GFP/+* |
| **H** | *eyFLP1/+; TubGal80, FRT40A/FRT40A, UAS-Eiger*$^{+W}$*; Act>y[+]>Gal4, UAS-GFP/ UAS-unc-13-RNAi (VDRC33609)* |
| **K** | *eyFLP1/+; Act>y[+]>Gal4, UAS-RFP/GFP-Wingless; FRT82B TubGal80/FRT82B* |
| **L** | *eyFLP1/+; Act>y[+]>Gal4, UAS-RFP/GFP-Wingless; FRT82B TubGal80/FRT82B, UAS-Eiger*$^{KB}$ |
| **N** | *eyFLP1/+; Act>y[+]>Gal4, UAS-RFP/+; FRT82B TubGal80/FRT82B* |
| **O** | *eyFLP1/+; Act>y[+]>Gal4, UAS-RFP/+; FRT82B TubGal80/FRT82B, UAS-Eiger*$^{KB}$ |
| **P** | *eyFLP1/+; Act>y[+]>Gal4, UAS-RFP/wg*$^{l-8}$*; FRT82B TubGal80/FRT82B, UAS-Eiger*$^{KB}$ |
| **Figure S1** | |
| **A** | |
| **Wild-type** | *nub-Gal4/+; UAS-CD8-PARP-Venus/+* |
| ***RpS3*/+** | *nub-Gal4/+; UAS-CD8-PARP-Venus, RpS3*$^{Plac92}$*/+* |
| ***RpS3*/+ +Puc** | *nub-Gal4/+; UAS-CD8-PARP-Venus, RpS3*$^{Plac92}$*/UAS-Puc* |
| **E** | *nub-Gal4, UAS-EGFP.CD63/RpL19* $^{K03704}$ |
| **G** | *w*$^{1118}$ |
| **H** | *RpS3*$^{Plac92}$*/+* |
| **Figure S2** | |
| **A** | *nub-Gal4/+; UAS-CD8-PARP-Venus, RpS3*$^{Plac92}$*/UAS-Puc* |
| **B** | *nub-Gal4/UAS-unc-13-RNAi (2999R-2); UAS-CD8-PARP-Venus, RpS3*$^{Plac92}$*/+* |
| **C** | *nub-Gal4/+; UAS-CD8-PARP-Venus, RpS3*$^{Plac92}$*/UAS-Rab3-RNAi (JF02020)* |
| **D** | *nub-Gal4/+; UAS-CD8-PARP-Venus, RpS3*$^{Plac92}$*/UAS-SNAP25-RNAi (JF02615)* |
| **E** | *nub-Gal4/+; UAS-CD8-PARP-Venus, RpS3*$^{Plac92}$*/UAS-Syt-RNAi (3139R-1)* |
| **F** | *nub-Gal4/+; UAS-CD8-PARP-Venus, RpS3*$^{Plac92}$*/UAS-Syt-RNAi (JF01461)* |
| **G** | *nub-Gal4/+; UAS-CD8-PARP-Venus, RpS3*$^{Plac92}$*/UAS-Rab27-RNAi (14791R-2)* |
| **H** | *nub-Gal4/+; UAS-CD8-PARP-Venus, RpS3*$^{Plac92}$*/UAS-Rab27-RNAi (JF02165)* |
| **I** | *nub-Gal4/+; UAS-CD8-PARP-Venus, RpS3*$^{Plac92}$*/UAS-ALiX-RNAi (HMS00298)* |
| **J** | *nub-Gal4/UAS-ALiX-RNAi (VDRC32049); UAS-CD8-PARP-Venus, RpS3*$^{Plac92}$*/+* |
| **K** | *nub-Gal4/+; UAS-CD8-PARP-Venus, RpS3*$^{Plac92}$*/* UAS-*Gαq-RNAi (17759R-2)* |
| **L** | *nub-Gal4/+; UAS-CD8-PARP-Venus, RpS3*$^{Plac92}$*/ UAS-Plc21C-RNAi (4574R-2)* |
| **M** | *nub-Gal4/ UAS-norpA-RNAi (VDRC105676); UAS-CD8-PARP-Venus, RpS3*$^{Plac92}$*/+* |
| **O** | |
| **+Puc** | *nub-Gal4/+; UAS-CD8-PARP-Venus/UAS-Puc* |
| **+unc-13-RNAi** | *nub-Gal4/+; UAS-CD8-PARP-Venus/UAS-unc-13-RNAi (VDRC33609)* |
| **+Rab3-RNAi** | *nub-Gal4/UAS-Rab3-RNAi (7576R-3); UAS-CD8-PARP-Venus/+* |
| **Figure S3** | |
| **A** | *nub-Gal4/ wg* $^{l-8}$*; UAS-CD8-PARP-Venus, RpS3*$^{Plac92}$*/+* |
| **C** | *nub-Gal4 /GFP-Wingless; UAS-CD63.mCherry/+* |
| **D and E** | *nub-Gal4 /GFP-Wingless; UAS-CD63.mCherry/RpS3*$^{Plac92}$ |
| **G and I** | *nub-Gal4, UAS-VHH4-CD8-HA/GFP-Wingless; UAS-CD63.mCherry/RpS3*$^{Plac92}$ |
| **H** | *nub-Gal4, UAS-VHH4-CD8-HA/GFP-Wingless; UAS-CD63.mCherry/+* |

**Table S5. Detailed Genotypes shown in Figures, Related to Figure 1-4 and S1-S3.**
